# Protein Language Model Performs Efficient Homology Detection

**DOI:** 10.1101/2022.03.10.483778

**Authors:** Mesih Kilinc, Kejue Jia, Robert L Jernigan

## Abstract

**Motivation:** There are now 225 million sequences in the UniProtKB database, as of January 2022 and 451 million protein sequences in the NCBI non-redundant database. This huge sequence data is ripe for analysis and can be extremely informative about biological function when analyzed with the appropriate methods. Evolutionary information such as the relationship among protein sequences is key to performing sequence analyses. Since sequence matching is one of the primary ways that annotations are found, higher-quality sequence matches yield a larger number of identified homologs. Thus, there is an essential need for a faster and more accurate homolog detection method to process the huge amount of rapidly growing biological sequences.

**Method:** Recently, we have seen major improvements in various predictive computational tasks such as structure prediction from the ever-improving artificial intelligence methods. One such approach has been to use language models to represent proteins numerically in a representation matrix (embeddings) while retaining context-dependent biochemical, biophysical, and evolutionary information. Computational transformer architectures that utilize attention neural networks can generate these context-aware numerical representations in an unsupervised fashion. One such use for these protein embeddings is remote homolog detection. In this work, we utilize protein language models and then apply discrete cosine transforms to extract the essential part of these embeddings, resulting in a significantly smaller fixed-size matrix for each sequence. This allows us to numerically and efficiently calculate the distance between all pairs of proteins resulting in homolog detection.

**Results:** Our Protein LAnguage model Search Tool (PLAST) is significantly faster, with linear runtimes in the number of sequences within the query database. With only one CPU core, it can scan a million sequences in less than a second. It essentially removes the noise in the sequence data and leads to significant improvements. PLAST is more accurate in the benchmarks tested from the PFAM, SCOP, and CATH databases than other approaches. When benchmarked with the PFAM database, the increase in the area under the receiver operating characteristic curve (AUROC) 3.1% when compared with NCBI-BLAST. The number of remote homologs that are detectable now is significantly larger and pushes sequence matches deeply into the usual twilight zone. Compared with the state-of-the-art profile-based homology search tools like CSBLAST, the increase was still 2.0%. PLAST can find remote homologs for a significant number of proteins that had been thought to be unique due to homolog detection failure. These homologs that are found usually have less than 20% sequence identity making them indistinguishable from noise with most other sequence matching methods.

**Conclusion:** PLAST is an accurate and fast homolog detection tool essential for easy and rapid progress to utilize the vast amount of data generated by next-generation sequencing methods. Quantization of sequence embeddings into highly-compressed noise-free representations with the use of direct cosine transforms allows for the efficient and accurate detection of normal homologs and remote ones that are undetectable by other sequence similarity methods. The PLAST web server is accessible from https://mesihk.github.io/plast.

## Introduction

As the cost of genome sequencing continuously goes down, more and more genomic data is being deposited. However, most of those data have not been analyzed. For example, SwissProt, a well-annotated database has only 565 thousand sequences. Analyzing this vast amount of data has become an important challenge in the field of bioinformatics. Some of the failures are well known where there are nearly identical 3D structures, but usual sequence matches fail. In a recent work, we have recently solved this problem and now obtain sequence matching closely corresponding to the structure matches (1). Evolutionary information that is used there is key to improved sequence analyses. Finding similar sequences is usually the first step in investigating any new sequence. Accordingly, homolog detection is one of the most used techniques to analyze sequences. And by far the most heavily used computational tool in biology. This is evident in the large number of BLAST citations (2).

Sequence similarity search first starts by pairwise scoring of sequences. These algorithms utilize substitution scoring matrices to score all of the amino acid substitutions throughout the alignment. The global alignment algorithm (3) and local alignment algorithm (4) are two exact solutions to the pairwise alignment problem. However, they are computationally expensive to perform on a large scale because of *O*(*n*^2^) time complexity. To speed up search times, successful heuristic-based methods developed such as BLAST (5) and FASTA (6). When sequence similarities between query and homolog proteins are high (>30%), these methods perform well. However, for so-called twilight zone proteins that have sequence identity 25-30%, these methods often fail to identify homologs (7–9). To remedy these problems, a number of variants of BLAST have been purposed. Profile alignment methods use multiple sequence alignments to generate a probabilistic model for the query protein. These profiles can then be used to perform a search on the target database.

Some frequently used examples of such method are PSIBLAST (10) and CS-BLAST (11) tools. Another direction of research has relied upon Hidden Markov Models (HMM). These methods can learn a probabilistic representation of a protein family and perform alignments based on HMM representations. Tools such as PHMMER (12), or HHSEARCH (13) are based on these HMM representations. Methods that utilize profiles can iteratively increase the quality of search by including newly found homologs to query profiled and repeat the search.

Despite the heavy use of sequence matching by biologists, homolog detection is far from a solved issue. Heuristics utilized in BLAST are known to cause homolog detection discrepancies when different E-Values are used, leading to confusion among users (14–17). Moreover, in a recent paper, Wisman et. all (18) proved that most genes that are thought to be lineage-specific are actually homolog detection failures. In the framework they presented, it is shown that current sequence similarity tools cannot distinguish rapidly evolving genes from random matches, resulting in homolog detection failure. This shows the limit for pairwise sequence similarity searches, and that these failures can drastically affect the interpretations of evolution.

In the meantime, machine learning methods have developed efficient ways to learn representations of large-scale data in ways that do not require expert input. This unsupervised learning has allowed researchers to utilize the vast volume of protein sequence data to train neural networks to generate context-aware numerical representations (embeddings) of amino acids in the input sequences. These representations retain biochemical, biophysical, and evolutionary information about the input sequences. Embeddings encode remote homology, protein family information, secondary structures, tertiary contacts, and mutational effects. When compared with current predictive bioinformatics tools, neural network-generated embeddings can usually either outperform state-of-the-art methods or reach a similar level of success (19).

The present work is based on the powerful neural network embeddings of protein sequences from Reference (19) and another recent paper (20) that utilizes inverse direct cosine transform (iDCT) quantization and dynamic time wrapping (DTW) to find protein similarity. Here we develop a new method named PLAST that utilizes iDCT on the embeddings from the ESM (Evolutionary Scale Modeling) protein language model (19). Quantization with iDCT results with a significantly more concise representation of sequences. Having smaller representations allows us to perform significantly more efficient exact searches without any heuristics. In a single-core CPU, PLAST can perform a search against a database of a million proteins in less than 520ms. PLAST can find remote homologs for proteins that previously had no related sequences. These same proteins have remote homologs that were not detectable by traditional sequence matching tools because of very low sequence identities. Our method is not only faster but also more accurate in application to the benchmark (21) that uses the Pfam (22), SUPERFAMILY (SCOPe 2.08) (3, 24), and CATH Gene3D (5, 26) datasets. PLAST is an alignment-free method that simply compares the embedding vectors using this accurate homology detection tool.

## Methods

The PLAST pipeline begins by obtaining the protein representation with the ESM1b protein language model. The ESM1b (meaning Evolutionary Scale Modeling, version 1b) is a transformer based neural network model inspired by the recent successes of transformers in the natural language processing field. Protein language modeling is similar to character-level language models since proteins use a limited vocabulary of 20 amino acids. The ESM1b model is trained by masking some of the residues from the input sequence and training the model to predict the masked residues. Hence the model must learn the connections between the residues. It is trained on Uniprac database from Uniprot (27) with 250 million sequences. The model is able to distinguish amino acid types by their biochemical properties. Although no evolutionary information is given as input, the model learns the family information of sequences. This ensures that the model understands the evolutionary relationships. The model can predict biophysical properties such as secondary structures and contacts between residues. Moreover, a variant of it can predict mutational effects (28). These results provide strong evidence that embeddings with the ESM1b protein language model can also be applied for homology detection. However, embeddings can contain noise in the sense of homolog detection since these same embeddings can be used to predict multiple properties. We address this issue by quantizing the embeddings to reduce the noise and get the essential information. When no quantization is applied and raw embeddings from ESM1b are used in homolog detection with the help of the dynamic time wrapping technique, we find that the accuracy in the sense of AUC is 7.4% lower compared to the quantized version, supporting the idea that the embeddings have more information than needed.

The inverse direct cosine transform (iDCT) quantization method was introduced recently (20). DCT is widely used for lossy compression resembling those used for the JPEG format. iDCT has some properties that are useful for sequence representation. First, it preserves the sequential nature of the embeddings. Secondly, after quantization, it maintains most of the input information.

For a protein with N residues the embedding will be in a 3D matrix having dimensions 34 × N × 1280, where 1280 is the embedding length of an amino acid and 34 is the number of output layers of the language model. We select a layer then reduce N, residue resolution, to 5 using iDCT quantization. This embedding reduces N × 1280 to 5 × 1280, for every protein. These dimension reduction methods are analogous to principal component analysis and removes some noise. This same type of dimensionality reduction iDCT can be applied again to the other dimension of the 5 × 1280 representation matrix to obtain a 5 × 44 matrix. Another compressed representation from different layer with size 3 × 85 is generated in the same way. Which layers to use and the final matrix sizes are selected by parameter optimization, which results in the smallest size with best accuracy. The intuition behind having two matrix representation for every protein is to have different levels of quantization with different layers to increase the representation capability. Finally, we have a 5 × 44 and 3 × 85 matrix representations for every query protein and every protein in the database. These representations are all precomputed and generated in advance. To compare two protein sequences, it is necessary only to take the sum of the differences of every element in the two set of representation matrices. For small values, the two proteins will be homologs. This reduction significantly speeds up the search operation by allowing PLAST to perform millions of protein searches in less than a second with a single-core CPU. The algorithm is highly parallelizable and uses CPU vector operations. A single protein is just represented by 475 bytes. The whole SwissProt database which contains around 565 thousand proteins requires only 250 MB of RAM which is actually smaller than the SwissProt FASTA file itself. The architecture of PLAST is shown in Fig. 1.

**Fig. 1.**
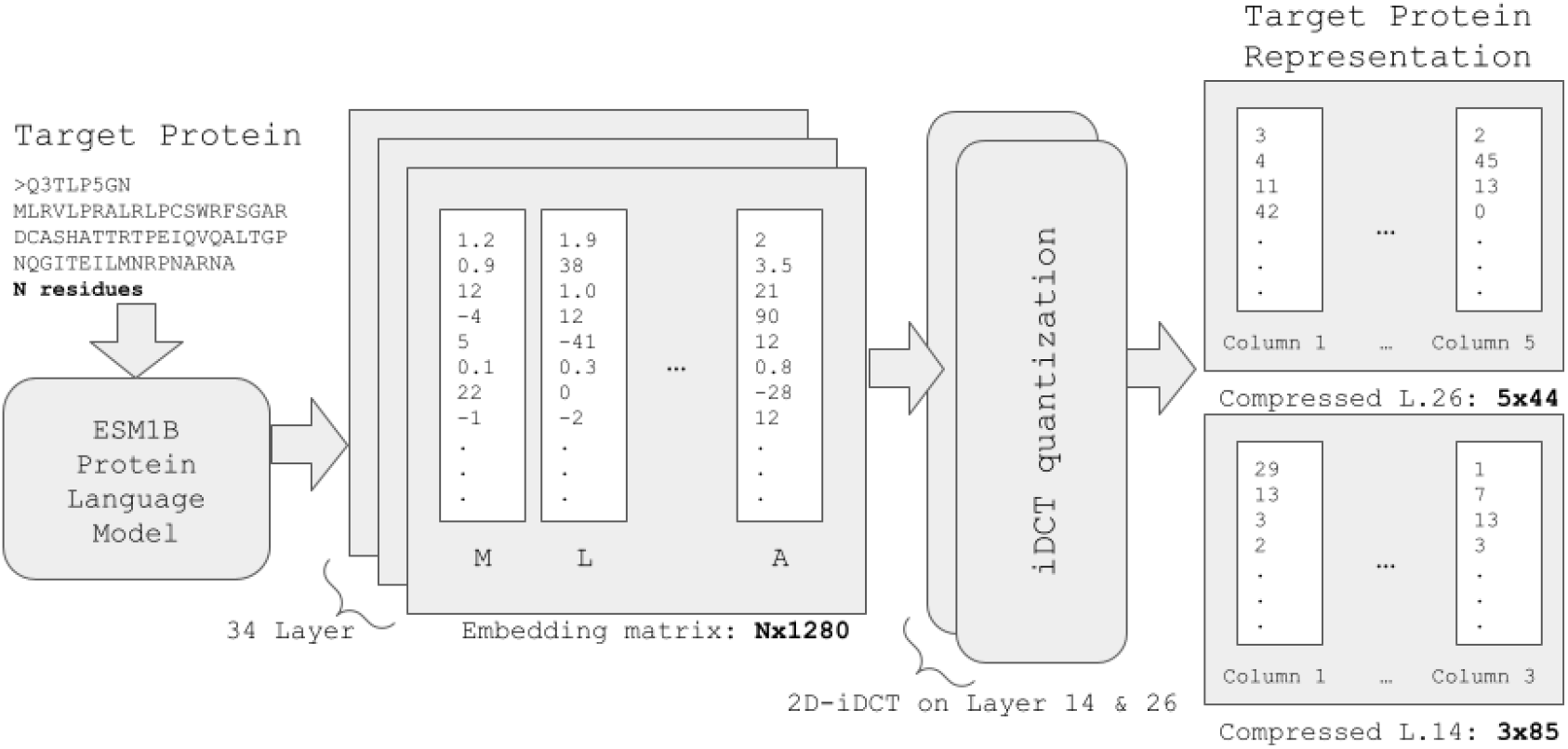
The architecture of PLAST. The first step is to feed the target protein to the ESM1b protein language model. This neural network model has 34 output layers and for each layer, it generates a vector of length 1280 for each residue. Accordingly, the output will be a 34 *×N ×* 1280 matrix where N is the length of the target protein. The next step is to compress the number of columns. The iDCT method is used to compress the layer-26 matrix from *N ×* 1280 to 5 *×* 1280. This is similar to moving from a residue representation to a domain representation. However, the compression ratio will be different for each individual protein since the protein size is not fixed in this representation. The next step is to apply the iDCT method to reduce it further from 5 *×* 1280 to 5 *×* 44 representation. Similarly, layer-14 compresses to 3 *×* 85. Finally, the target protein is represented with two fixed 5 *×* 44 and 3 *×* 85 matrices.

In the evaluation of our method, we use a recent benchmark (21). It contains three different datasets from Pfam (22), Gene3d (25) and SUPERFAMILY (23). These datasets contain proteins with multiple domains. If two proteins have the same domains, they are indicated to be homologs. If they don’t share any domain in common then they are annotated to be non-homologous. The Pfam benchmarking dataset contains 5,245 homolog pairs. The Gene3d dataset has 5,047 pairs. The SUPERFAMILY contains 5,656 pairs. Every dataset also contains the same number of non-homolog pairs as homolog pairs. The total number of protein pairs that are considered in the benchmark is 31,896. We compared the PLAST accuracy with the same metrics represented in benchmarking paper (21). The area under the ROC curve (AUC) and AUC score calculated on the first 1000 false positives AUC1000 are taken as the metrics for performance.

## Results

PLAST uses a compressed representation scheme for proteins. Both query protein and target database proteins are represented with only 475 bytes. This is achieved by utilizing the ESM1b protein language model embeddings and quantizing them with inverse direct cosine transforms in both dimensions to obtain the essence of the embedding. This allows us to have noise-free, minimized domain signatures for proteins under scrutiny while having the best accuracy compared with three different datasets

There are several objectives in the choice for the protein representation. (1) Removing the noise from the embeddings. Surprisingly, when no compression is applied in both of the dimensions (Layer 34 *N* × 1280), the accuracy for the Pfam dataset given as the AUC metric is already 91.6%, only 7.4% lower than the selected compression scheme (Layer 26 5 × 44 and layer 14 3 × 85). (2) Reducing the size of the database, so it will fit into RAM memory, for faster search times. Our goal is to fit the NCBI non-redundant database into commodity server-grade memory. (3) A small number of columns in the protein representation matrix won’t need alignment so that the distance will be sufficient for searching and scoring as explained below. The AUC score for the Pfam dataset is used to select the best parameters for the compression. Fig. 2-a shows the heatmap of AUC scores with different parameters selections with the distance metric. It can be seen from the results that as the number of columns increases, the distance metric performs poorly since we are nearer to a residue representation. However, a higher row count increases the AUC score as more information is available for comparison. Nevertheless, high dimensionality increases the demand for memory, resulting in database fragmentation leading to substantially longer search times. Fig. 2-b shows the difference in Pfam AUC scores when the dynamic time wrapping algorithm (alignment) and distance metric comparison is used. It can be seen that alignments have increased AUC scores. However, alignment needs prohibitively more time. Fig. 2-c shows the performance for different layers of ESM1b. The first layers perform poorly. However, the best performing layer is not the last layer (34) instead it is layer 26. In the optimization studies, we found that using two different layers with two different compression levels increases the performance considerably. Finally, a compression scheme that uses layer 26 embeddings with 5 columns and 44 rows and layer 14 embeddings with 3 columns and 85 rows is the best option to have state-of-the-art accuracy without requiring actual alignment. The memory footprint for each protein is a humble 475 bytes resulting in easily manageable database sizes. We have used different distance measurement methods for protein representation comparison such as L1 (eq. 1), L2 (eq. 2,) inner product distances where inputs linearized before distance calculation, dynamic time wrapping algorithm (DTW), Hausdorff method, and Frobenius norm. Here, DTW is a dynamic programming algorithm for calculating the similarity between two matrices. It is similar to the global alignment algorithm but applied to more broad problems such as speech recognition, signature recognition, etc. In this study, we used a fast implementation of DTW (29) that has linear *O*(*N*) runtime.

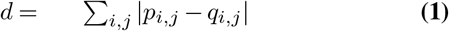

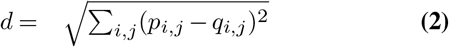

**Fig. 2.**
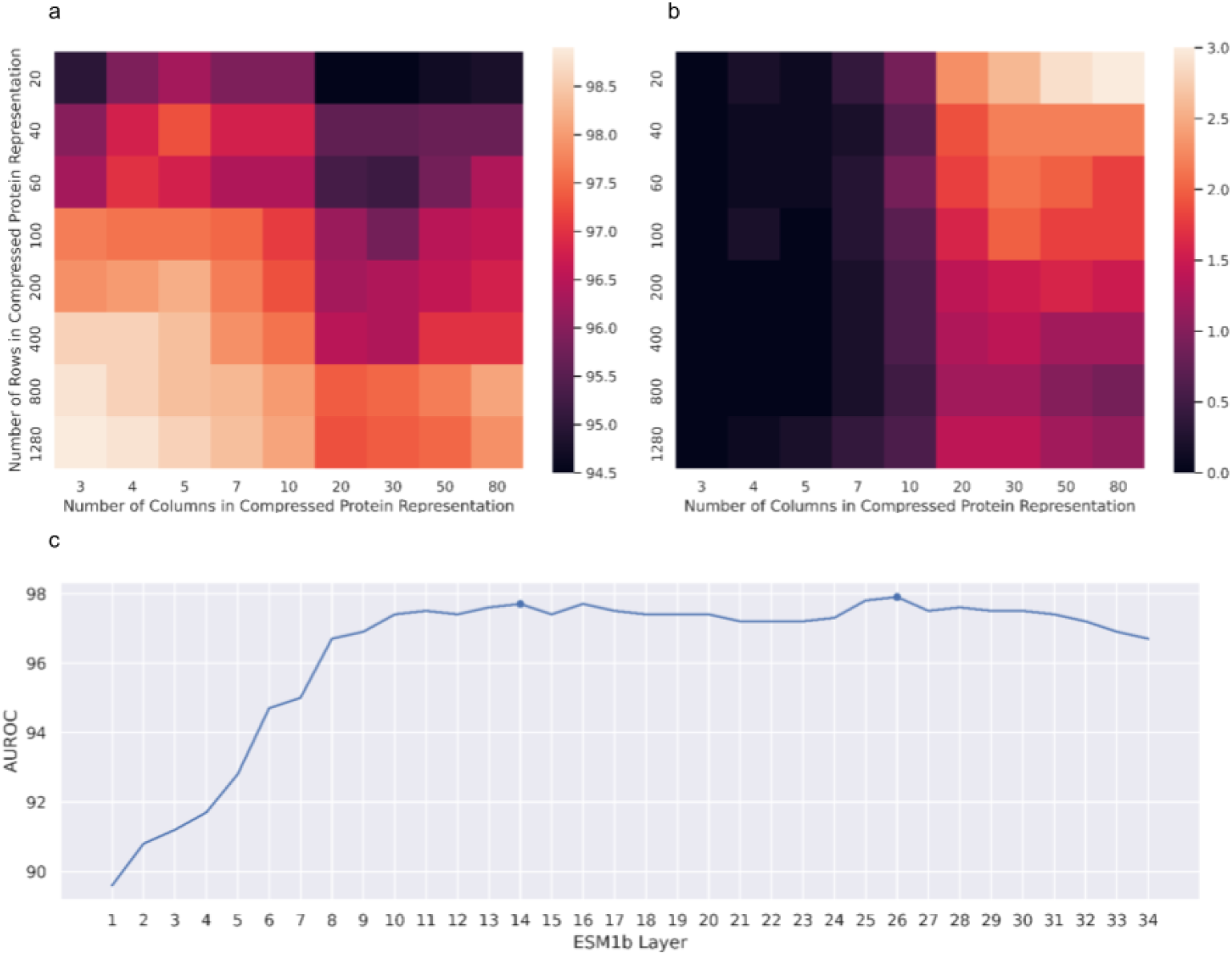
Results of Different Levels of Compression. (a) Protein representation matrix column and row count selection effect on the Pfam AUC scores when the distance metric is used. A grid search is performed to find the best compression ratio for protein representations. This figure represents the AUC score of the Pfam benchmark dataset when different column and row sizes are used for quantization. The distance metric has good prediction capabilities when the number of columns is < 10, and for such low dimensionality alignments won’t have much impact. As the information content increases, we see an increase in the AUC scores. 5 *×* 44 is selected because it maximizes the AUC score while minimizing the amount of storage needed. (b) The difference in the AUC scores between the distance and the DTW technique (global alignment). When the column dimensionality is below 10, the differences are negligible. The distance calculation has a time complexity advantage compared to the DTW method. (c) Overall prediction performance over whole benchmarking dataset when different layers of ESM1b protein language model are used with 5 *×* 44 quantization. Layer 26 has the best prediction capability. Layer 14 is coupled with layer 26 to increase performance on PLAST-S. These layers are indicated in the plot.

**Fig. 3.**
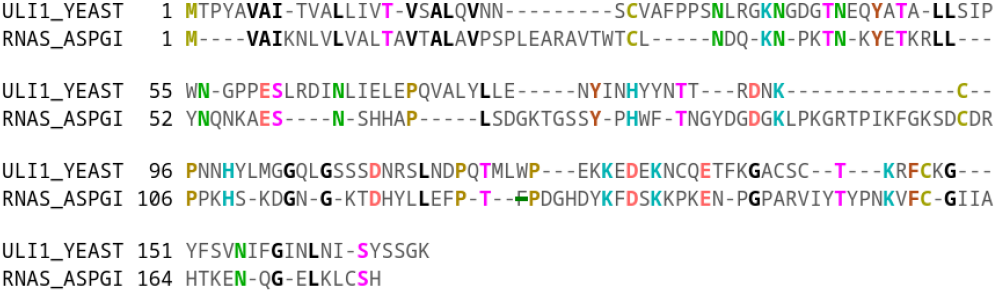
Alignment of ULI1 and homolog P00655 identified with PLAST. In this alignment, the ProtSub substitution matrix with gap opening penalty 5 and gap extension penalty 1 is used. Sequence identity was **25.2%** with a gap ratio 41.3%. The alignment score was 154. Identical residues are colored based on chemical features. This alignment is visualized using the bioinformatics.org/strap/aa tool.

**Fig. 4.**
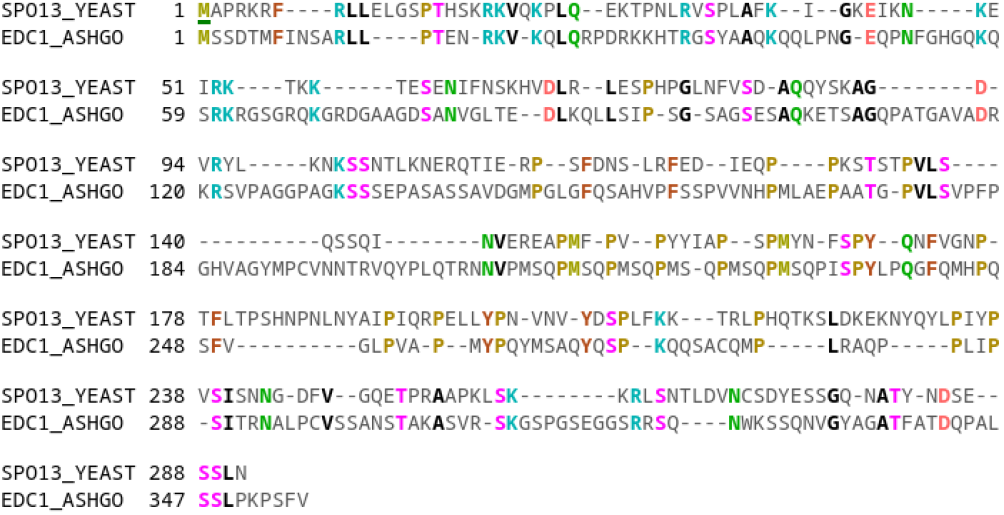
Alignment of SPO13 and homolog Q753U4 found by PLAST. In this alignment, the ProtSub substitution matrix with gap opening penalty 5 gap and extension penalty 1 is used. Sequence identity is found to be **24.3%** with a 38.1% gap ratio. The alignment score is 259. Identical residues are colored based on chemical features. This alignment is visualized using the bioinformatics.org/strap/aa tool.

The best distance metric is the dynamic time wrapping technique because it is similar to global alignment. However, as in the global alignment case, this technique also increases time complexity even though there are efficient implementations such as fastdtw (29). The second-best distance method is the L1 distance. For low dimensionality (<10) the L1 and DTW metrics perform similarly. The differences for the Pfam dataset AUC score with L1 and DTW distances are shown in Fig. 2-b. When the number of columns is ≤ 5 then there is no difference in the sense of accuracy between the L1 and DTW technique since for such low dimensionalities, alignment does not add any benefit. But, when there are more than 10 columns and information content is low then the DTW method performs much better. However, thanks to modern

CPU architectures, vector-based operations can be done in one CPU cycle with SIMD extensions. As a result, the distance calculation can be performed in constant *O*(1) time. This is much better compared to *O*(*n*^2^) time required for the DTW method. Accordingly, a comparison of a pair of proteins can be done in constant time with the distance metric.

The validation of the PLAST tool has been carried with the most commonly used alignment-based homolog detection tools. PFAM, GENE3D, and SUPERFAMILY datasets were used in the validation efforts. AUC and AUC1000 scores, the AUC score calculated for the first 1000 false positives andindicates the quality of the ranking, are used to show the performance of the tools as shown in Table 1. We have compared with the CSBLAST (11), PHMMER (12), HHSEARCH (13), NCBI-BLAST (5), UBLAST (30), and FASTA (6) scores reported in the detailed benchmarking paper (21). Moreover, we also compared with the method used in the ESM1b paper (19). In this method, they calculated the mean of the last (34th) layer of ESM1b embedding columns to have a fixed size representation of a protein. This result is shown as ESM1bL34M. Table 1 shows how PLAST performs better in every case compared to these other tools.

**Table 1.**
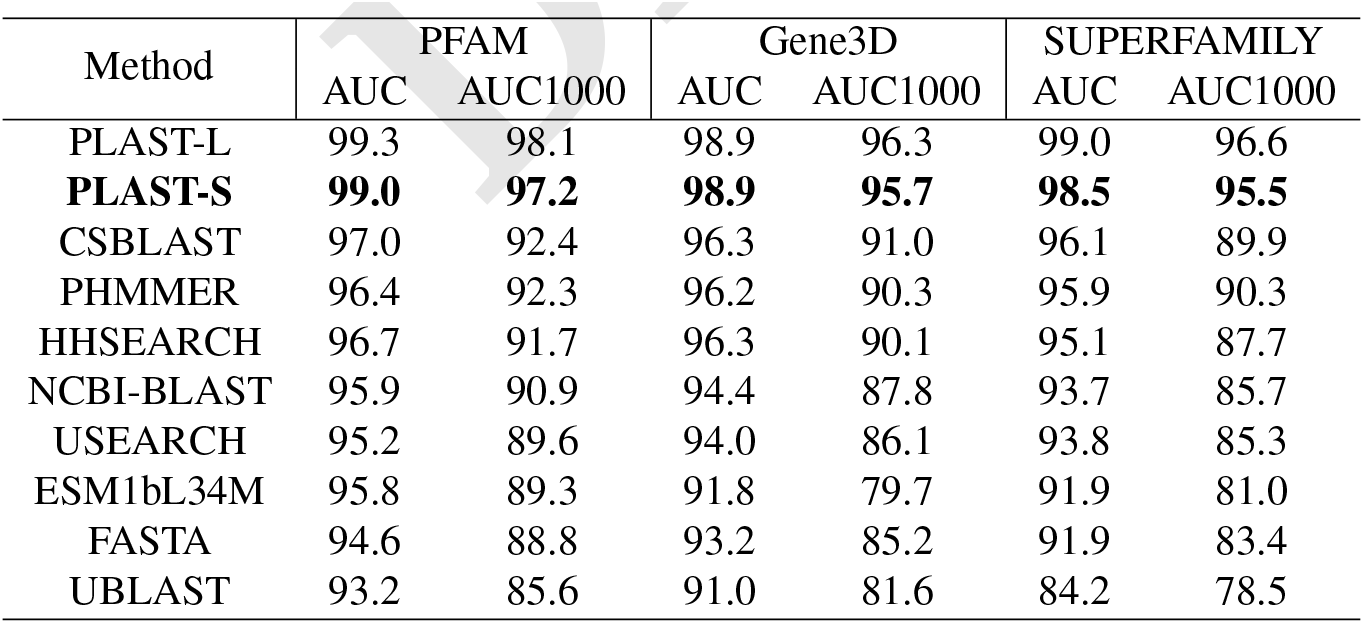
Comparison of PLAST performance on the PFAM dataset

Firstly, we note that compression parameters of PLAST have been selected to minimize the storage requirement while performing better than state-of-the-art methods. However, with a more complete representation it is possible to achieve even higher results. Table 1 shows these results as PLAST-L and compact representations as PLAST-S. PLAST-L uses a compression scheme as layer 14 with 3×1280 and layer 26 with 3×1280 totaling 7680 bytes per protein while PLAST-S only uses 475 bytes.

Table 1 shows the performance of commonly used tools when benchmarked on the Pfam database. It shows that PLAST is the best tool with both the AUC and the AUC1000 metrics. PLAST in comparison with the closest tool, CS-BLAST, has a 2% higher AUC score and a 4.8% higher AUC1000 score. Compared to the commonly used NCBIBLAST tool, PLAST has 3.1% higher AUC and 6.3% higher AUC1000 scores. When benchmarking is performed with the Gene3D database, PLAST outperforms CS-BLAST with a 2.6% higher AUC score and 4.7% higher AUC1000 score. Lastly, in the SUPERFAMILY dataset, PLAST is the best tool with the 2% higher AUC score than CS-BLAST, and in terms of AUC1000, a 5.2% higher score compared with PHMMER (PHMMER has a higher AUC1000 score than CS-BLAST).

These results strongly suggest that PLAST is an accurate homology detection tool outperforming commonly used alignment-based homolog detection tools for the Pfam, Gene3D, and SUPERFAMILY datasets while minimal possible representation. These consistently permit homolog identification at lower levels of sequence identity.

Benchmarking results can be used to select a threshold for the distance metric to identify homologs and non-homologs. Using the benchmarking dataset, we have calculated *F*_1_ scores based on (eq. 3) for different thresholds and selected 6828.5 since it maximizes the *F*_1_ score.

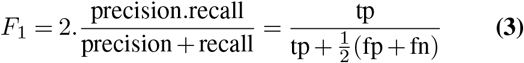

PLAST databases have a very small memory requirement due to the space-efficient protein representations. A protein sequence in a PLAST database is represented with only 475 bytes of data. This allows us to store the whole database in memory, which leads to significantly faster search operations. The database search in PLAST actually performs distance calculations between representation vectors. Thanks to the current CPUs that have vector-enabled instruction sets, PLAST can perform an exact search for millions of proteins in less than a second with only a single core. Table 2 shows the memory requirements and exact search times for popular and commonly used databases when only a single-core AMD Ryzen 4800H is used. The SwissProt database requires only 250MB of memory which is smaller than the SwissProt FASTA file itself. The exact search time for SwissProt is just 285ms. On the other hand, the NCBI non-redundant (nr) protein sequence database which is one of the largest databases requires only a 198.88GB of memory, an amount of memory that is commonly available in modern servers. An exact search on the full NCBI non-redundant database can be performed within 225.72 seconds. Note that these times are for a single-core CPU, as an example, if 32 core parallelization were utilized, then the search time for the entire NCBI non-redundant database is only 7.05 seconds. These memory and timing results are extrapolated based on actual one-core Swissprot results.

**Table 2.** PLAST-S memory requirement and search time for commonly used databases

| Database | Num. of Seq. | Memory | Search Time | 32-Core Parallelization |
| --- | --- | --- | --- | --- |
| Swissprot | 565,928 | 0.25G | 0.28s | 8.9ms |
| TrEmbl | 225,013,025 | 99.54G | 112.97s | 3.53s |
| Uniref100 | 280,483,851 | 124.08G | 140.82s | 4.40s |
| NCBI-Non Redundant | 449,569,698 | 198.88G | 225.72s | 7.05s |

## Discussion

The method of homolog detection in PLAST is radically different compared to traditional sequence or HMM profile alignment tools. We investigated the effects of this difference on the ability to identify homologs as well as the performance with different metrics. In a recently published paper, it is shown that as the rate of protein evolution increases, the probability of remote homolog detection decreases with traditional sequence similarity search tools. This happens because the sequences become so divergent that the tools can’t distinguish them from random matches. The mathematical framework laid by Weismann et. all (18) can predict whether the rate of evolution is too high for remote homolog detection with traditional sequence alignment-based tools. In their study, they found that most genes that don’t have any homologs outside of specific species (lineage-specific genes) are wrongly attributed to biological novelty. They presented that the majority of lineage-specific genes are actually the result of homolog detection failure. If a more sensitive synteny analysis were performed it was shown that 46% of genes are found to have homologs (18). Motivated by these results we performed a PLAST search on two *Saccharomyces cerevisiae* proteins, ULI1 and SPO13, and found remote homologs from (P00655) *Aspergillus giganteus* and (Q753U4) *Ashbya gossypii*. Those proteins were previously thought to be novel because of homolog detection failures. But PLAST appears to be better at finding such remote homologs.

Another example is HPO30 protein, an uncharacterized protein of Caenorhabditis elegans. NCBI-BLAST and PHMMER are not able to find any homologous sequence to HPO30 in the SwissProt database. However, it is known that this protein belongs to PFAM claudin-like domain (PF07062). PLAST readily identifies a hundred of homologs to HPO30 that have around 20% sequence identity. One example alignment with human homolog CLRN3 (Q8NCR9) is shown in Fig. 5-a. HPO30 and CLRN3 have 21.5% sequence identity however, they have the same structure fold. Structural alignment of Alphafold2 predictions of these proteins are given in Fig. 5-b (31). Overall core residue alignment has an RSMD of 3.4 Å.

**Fig. 5.**
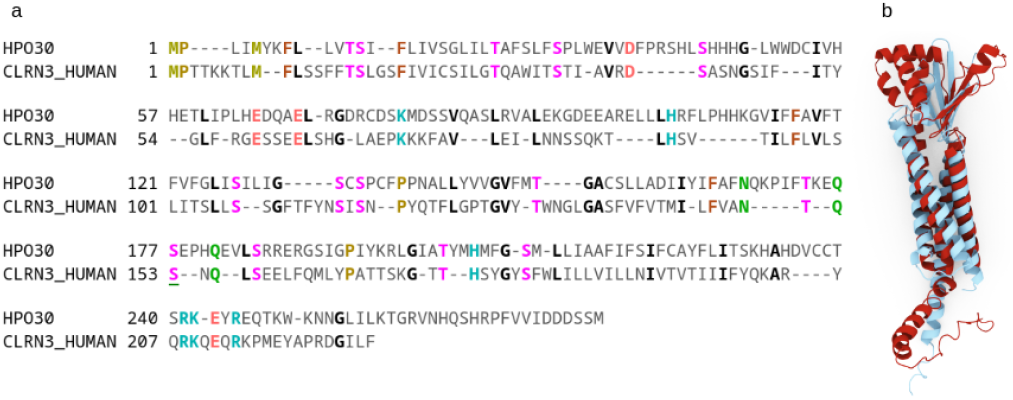
Detection of a Remote Homolog for HPO30 and Validation with Predicted Structure Comparison. (a) Alignment of HPO30 and human homolog Q8NCR9 found by PLAST. In this alignment, the ProtSub substitution matrix with gap opening penalty 5 gap and extension penalty 1 is used. Sequence identity is **21.5%**, with gap ratio of 32.8%. The alignment score is 189. Identical residues are colored based on chemical features. This alignment is visualized using bioinformatics.org/strap/aa tool. (b) Structure matching of the two Alphafold2 (31) predicted structures of HPO30 (red) and CLRN3 (blue) human homolog that is found by PLAST. Structural alignment of 160 core residues has an RSMD of 3.4 Å.

One more experiment was performed for HIV gagPol polyprotein (P04585) to assess the performance of PLAST for an easy homolog detection task. For this protein BLAST found 304 homologs on SwissProt. PLAST found 325 homologs. 115 of them were identical in both results. This shows that PLAST can be used not only for remote homolog but also for normal homolog detection.

PLAST is a global homology tool. PLAST uses a fixed-length (variable-ratio quantization) representation of proteins. Combined with simple distance calculations this allows the fastest search operation possible. However, this feature undermines the ability to detect local homology. A local fragment for a homolog will not usually be detected by PLAST. Fig. 6 shows the predicted outcome of protein pairs with the differences in sizes between query and target proteins. It can be seen that as the size difference increases PLAST performance decreases. PLAST is clearly better at whole protein homolog identification.

**Fig. 6.**
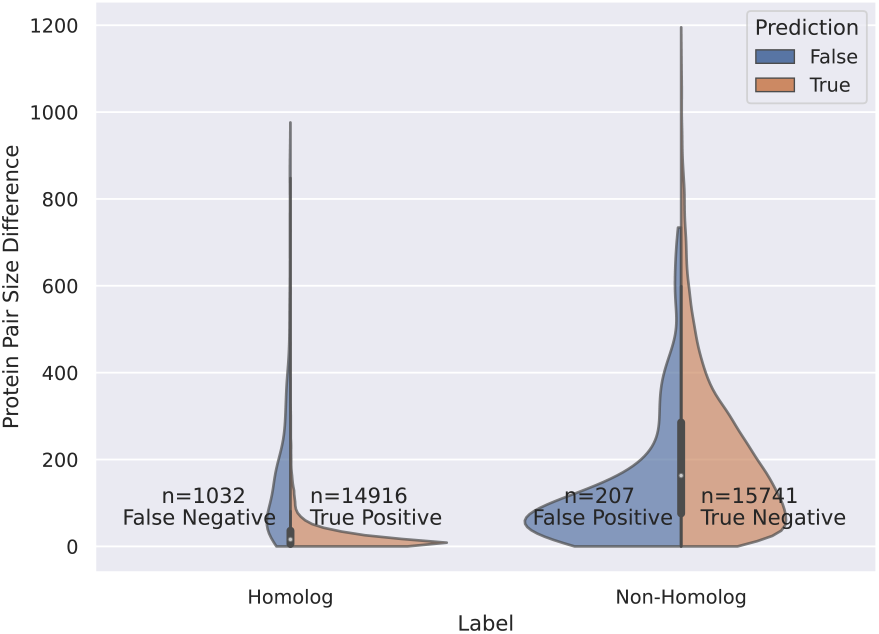
Prediction outcomes of benchmark protein pairs shown for different protein size differences in the number of residues between two proteins being compared. For example, if a pair of proteins have lengths of 120 and 230 residues are compared, then the difference in length will be 110. Most protein pairs in the benchmarks have similar lengths. This figure shows the distribution of predicted results based on the homology label when a threshold of 6828.5 is used (see ‘threshold’ selection above). This threshold is based on the benchmarking dataset found by the maximized F1 score.

## Conclusions

In this paper, we present PLAST, an exact search-based extremely fast, and accurate homolog detection tool that utilizes protein language models. It is a radically different method from other search methods in being an alignment-free and sequence similarity-free architecture that uses reduced protein representations. PLAST outperforms commonly used traditional alignment-based tools for benchmarks from major databases such as PFAM, Gene3d, and SUPERFAMILY. PLAST is memory efficient and can keep hundreds of millions of protein representations in the memory of a server-grade computer. PLAST is not limited to theoretical limits of sequence similarity-based tools. It can find remote homologs for proteins that have a high evolution rate that makes them indistinguishable from noise with traditional alignment-based tools, and these results push sequence alignment deep into the twilight zone.

## ACKNOWLEDGEMENTS

We are grateful to the NIH for support from Grant R01GM127701 and Ricardo Henriques for his BioRxiv LATEX template.

